# Molecular and cellular dissection of the OSBP cycle through a fluorescent inhibitor

**DOI:** 10.1101/844548

**Authors:** Tiphaine Péresse, David Kovacs, Mélody Subra, Joëlle Bigay, Meng-Chen Tsai, Joël Polidori, Romain Gautier, Sandy Desrat, Lucile Fleuriot, Delphine Debayle, Marc Litaudon, Van-Cuong Pham, Jérôme Bignon, Bruno Antonny, Fanny Roussi, Bruno Mesmin

## Abstract

ORPphilins, natural molecules that strongly and selectively inhibit the growth of some cancer cell lines, are proposed to target intracellular lipid-transfer proteins of the Oxysterol-binding protein (OSBP) family. These conserved proteins exchange key lipids, such as cholesterol and phopsphatidylinositol-4-phosphate (PI(4)P), between organelle membranes. Among ORPphilins, molecules of the schweinfurthin family interfere with intracellular lipid distribution and metabolism, but their functioning at the molecular level is poorly understood. We report here that cell line sensitivity to schweinfurthin G (SWG) is inversely proportional to cellular level of OSBP. By taking advantage of the intrinsic fluorescence of SWG, we follow its fate in cell cultures and show that its incorporation at the TGN depends on OSBP cellular abundance. We report that SWG inhibits specifically the lipid exchange cycle of OSBP. As a consequence, post-Golgi trafficking, membrane cholesterol levels and PI(4)P turnover are affected. Finally, we demonstrate the direct binding of SWG into OSBP lipid-binding cavity by intermolecular FRET. Collectively these data describe for the first time a specific and intrinsically fluorescent pharmacological tool to dissect OSBP properties at the cellular and molecular levels.

## Introduction

ORPphilins are small natural bioactive molecules that have strong inhibitory activities on the growth of several human cancer cell lines. Despite their structural diversity, these compounds may have a related mechanism of action because they present a similar sensitivity profile on the NCI-60 cancer cell panel (1). It is proposed that ORPphilins directly target Oxysterol-binding protein (OSBP) and OSBP-related protein 4 (ORP4), which are two members of a conserved family of lipid-transfer proteins (LTPs) that transport key lipids such as cholesterol or phosphatidylserine between intracellular membranes (2, 3). OSBP is a cytosolic protein containing an N-terminal region capable of bridging the membrane of the endoplasmic reticulum (ER) to that of the trans-Golgi network (TGN), and a C-terminal OSBP-related domain (ORD) responsible for the lipid transfer activity (4). We have previously demonstrated that OSBP directs cholesterol transport from the ER to the TGN through coupled counter-exchange and hydrolysis of the phosphoinositide PI(4)P at membrane contact sites between these two organelles (5). By using the ORPphilin OSW-1 on cells whose endogenous level of OSBP exceeds that of ORP4 by two orders of magnitude, we have shown that OSBP activity makes a major contribution to the intracellular trafficking of cholesterol (6). As a result, OSBP impacts on the physicochemical properties of cellular membranes. The other potential target of ORPphilins, ORP4, is the closest OSBP ortholog and is also described as a cholesterol/PI(4)P exchanger (7), among other assigned roles (8). However, ORP4 is virtually absent from most tissues except the brain, testis and some transformed tissues, while OSBP is ubiquitously expressed.

Among the ORPphilins described by Burgett *et al.* are the small molecules schweinfurthin A and B, which were shown to compete with oxysterol binding to OSBP and ORP4 (1). Interestingly, however, unlike other ORPphilins, the affinity of these compounds was 40 times higher for OSBP than for ORP4, suggesting that schweinfurthins have a better binding specificity to OSBP (1). It has also been shown that cell treatment with schweinfurthin A could shift OSBP localization to the Golgi area (1). The compounds from the schweinfurthin series (including vedelianin and schweinfurthins A-Q) are natural prenylated stilbenes isolated from the leaves or fruits of different species of the tropical genus *Macaranga* (9–14). Schweinfurthins harboring a hexahydroxanthene moiety are promising because they have a powerful and selective antiproliferative activity against some human cancer cell lines (10, 15). For example, schweinfurthins show GI_50_ values in the nanomolar range on cell lines derived from glioblastomas, while they have only limited activity on lung cancer cell lines, such as A549 (13, 16). However, this differential cytotoxicity is still poorly understood (15). Previous studies on the mechanism of action of these drugs have shown that they interfere with intracellular trafficking and lipid metabolism. For instance, the compound 3-deoxyschweinfurthin B reduced cellular cholesterol levels by decreasing its synthesis and increasing its export (17). Likewise, the treatment with schweinfurthin G (hereafter abbreviated as SWG) resulted in the disruption of the TGN and the depletion from the cell surface of some lipids (GM1, sphingomyelin and cholesterol) known to promote membrane order and domain formation (18). In these conditions, PI3K-Akt-mTOR signaling pathway was downregulated (18).

At the molecular level, do schweinfurthins bind directly to OSBP? A biotinylated analogue of SWG has been shown to bind to OSBP, ORP1L and ORP1S, albeit with low affinities (in the μM range) (18), which contrasted with the nanomolar concentrations needed to observe a cellular effect. More surprisingly, a bioactive fluorescently-tagged SWG showed a punctate intracellular distribution (18, 19), partially localized with an endosomal marker, but far from the ER/TGN interface where OSBP concentrates. Nevertheless, an interesting relationship between schweinfurthins and OSBP has been established by the fact that OSBP knockdown sensitizes cells to schweinfurthins, and conversely, that OSBP overexpression increases their resistance to schweinfurthins (1).

Altogether, these previous observations suggest that schweinfurthins may have an inhibitory effect on OSBP, but whether this effect is direct or indirect is not well characterized. Here we dissect the fine interaction of the highly active compound SWG with OSBP using in vitro reconstitution of membrane systems and cellular imaging approaches. We reveal and take advantage of the fact that SWG has intrinsic fluorescence properties. This allows us first to track SWG within cells with good spatial and temporal precision, and second to perform OSBP binding kinetics. We report the nanomolar affinity of SWG for OSBP, its sensitivity to membrane interfacial environment, and provide evidence that SWG is able to inhibit the entire lipid exchange cycle catalyzed by OSBP by entering its ORD domain.

## Results

### OSBP expression level correlates with SWG sensitivity

The specific cytotoxicity of schweinfurthins for a subset of NCI-60 cancer cell lines is not explained so far. Understanding what differentiates cell lines that do not respond identically to SWG would allow a more detailed molecular analysis of its activity (**Fig 1A**). Previous data have shown that modulating the cellular level of OSBP (via knockdown or overexpression) could influence the sensitivity of cells to schweinfurthins (1). We therefore thought that the differential cytotoxicity to schweinfurthins observed between various cell lines could be due to their divergent levels of OSBP expression. We then compared OSBP expression in a panel of several cancer cell lines, to which we added the immortalized retinal pigmented epithelial cell line (hTERT-RPE1; hereafter RPE1) that we previously identified as sensitive for OSW-1 (6), and measured their sensitivity to SWG. We found that OSBP level varies up to 5 fold between these cell lines (**Fig 1B**). OSBP expression is relatively low in human primary glioblastoma cell line U87 MG, as well as in RPE1, while it is higher in PC3 (human prostate cancer cell line) and maximal in A549 cells (adenocarcinomic human alveolar basal epithelial cells). We then performed viability tests after treatments with increasing doses of SWG for 72 h, and plotted the half maximal inhibitory concentrations (IC_50_) obtained for each cell line according to their OSBP level (**Fig 1C**). We observed that the sensitivity to SWG is inversely proportional to the cellular level of OSBP, that is, high expression correlated with resistance. As a control, we used Doxorubicin, a compound not related to ORPphilins known to target DNA metabolism. In this case, we do not observe any correlation between OSBP expression level and Doxorubicin sensitivity (**Fig 1D**), and confirmed that the sensitivity profile for Doxorubicin is different from that of SWG (**Fig 1E**). These results indicate that OSBP has a protective effect against the cytotoxic activity of SWG. Note that a good correlation between sensitivity/resistance and expression level generally defines drugs with a single protein target (20).

**Figure 1.**
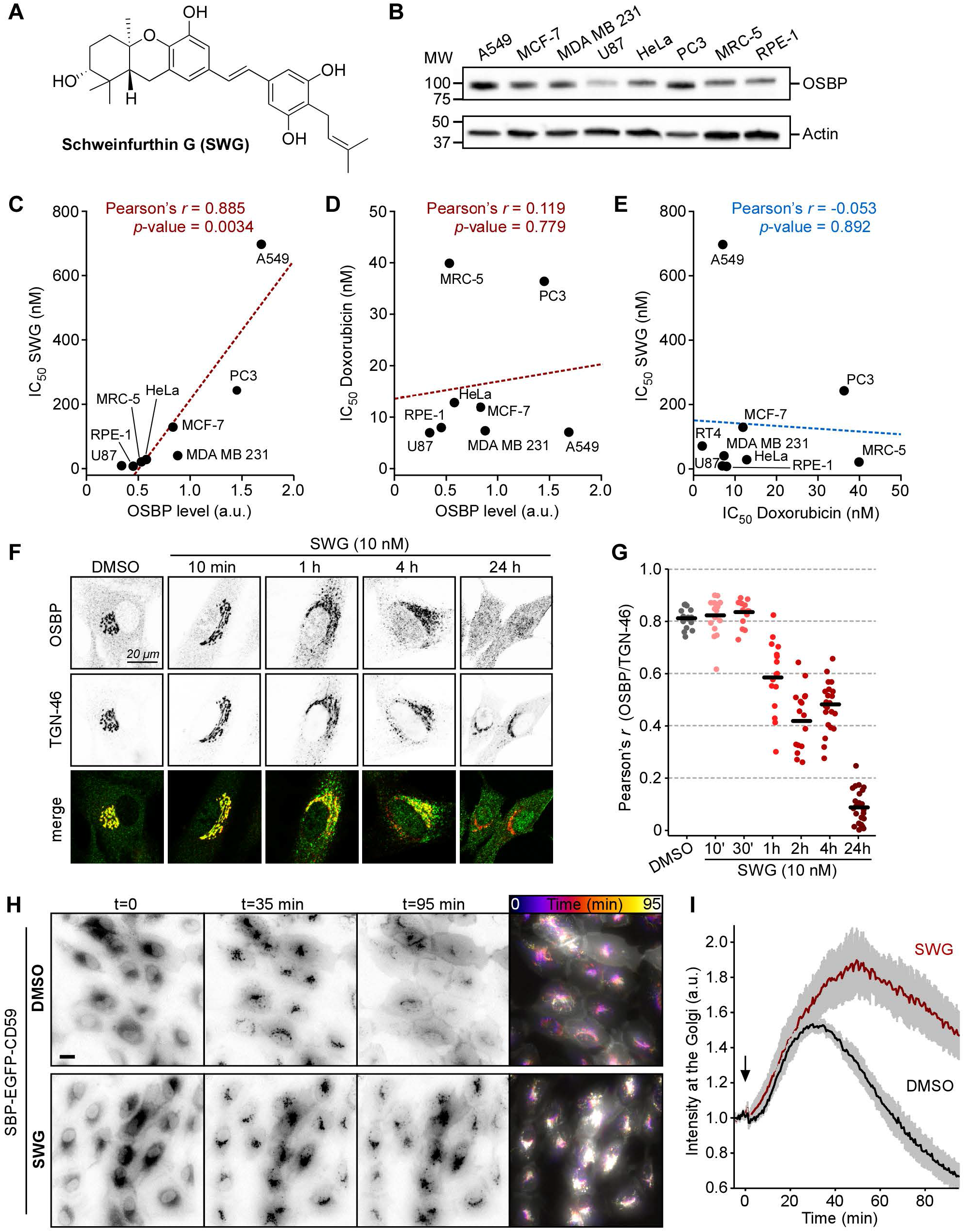
Characterization of SWG sensitivity and effects on TGN trafficking. **A.** Structure of Schweinfurthin G (SWG). **B.** The immunoblot shows the relative level of endogenous OSBP expression in the indicated cell lines. **C, D.** Measurements of cell viability (IC50 values) for the different cell lines treated with SWG (C) or Doxorubicin (D) are plotted as a function of the normalized OSBP expression level found in each cell line. **E.** The sensitivity profile for SWG is plotted as a function of that for Doxorubicin for each cell line used. The indicated Pearson’s correlation values (*r*) and *p*-values were determined using SigmaPlot. **F.** RPE-1 cells cultured for 24 h were treated with DMSO or SWG as indicated. Cells were then fixed, permeabilized, and processed for immunofluorescence to assess the localization of endogenous OSBP and TGN-46. Confocal microscopy images are single optical sections. **G.** Evolution of the Pearson’s correlation coefficient between OSBP and TGN-46 upon SWG treatment over time, or DMSO as control. Measurements were performed on 15–25 cells for each condition, from 3 independent experiments with the mean values indicated by the black lines. **H.** Time-lapse imaging of RPE-1 cells stably expressing SBP-EGFP-CD59 incubated with biotin and, as indicated, DMSO or SWG (50 nM), for the indicated time. Real-time images were acquired using a wide-field microscope at 2 frames/min. Temporal projections (right image) were performed on entire stacks of images. **I.** Normalized intensity of SBP-EGFP-CD59 at the Golgi. Data are means ±SEM (n=4). Scale bars: 20 μm.

### SWG modulates TGN-to-PM trafficking

Because OSBP is most likely the target of SWG, we next wanted to monitor OSBP intracellular distribution over time after adding SWG. We used RPE-1 cells because they are highly sensitive to SWG and have a wide and extensively tubulated TGN that is well adapted to imaging. **Fig 1F and** **1G** show that in control cells, endogenous OSBP is predominantly co-localized with the TGN marker TGN-46, and weakly present in the cytosol. Upon short treatments with low doses of SWG, we noticed that this cytosolic fraction of OSBP shifted completely to the TGN. In contrast, more prolonged treatments with SWG lead to the gradual dissociation of OSBP from the TGN, as judged by Pearson’s correlation coefficient measurements over time (**Fig 1G**), until it became mostly cytosolic after 24 hours of treatment. In agreement with previous observations, long treatments with SWG also lead to TGN disruption (18). In particular, the TGN loses its characteristic tubular morphology and becomes partly dispersed. These short- and long-term effects, first on OSBP distribution and then on TGN morphology, suggest that SWG could quickly and durably affect Golgi trafficking.

To further analyze the effect of SWG on the secretory pathway, we sought to monitor the intracellular transport of a newly synthesized protein from the ER to the plasma membrane via the Golgi. We therefore used the RUSH system, which allows to trigger the secretion of a cargo (here, the CD59 protein) fused with a streptavidin binding peptide (SBP) by simply adding biotin to the culture medium (21). Since CD59 is also coupled to GFP, we could track and quantify its trafficking by live cell imaging. Experiments performed with control cells at 37°C in DMEM/F12 medium showed that CD59 initially retained in the ER (t=0) crosses the Golgi apparatus within 35 min after biotin addition, and finally reaches the plasma membrane after 90 min (**Fig 1H**). **Fig 1I** illustrates this wave of secretion by showing the fluorescence intensity in the Golgi region at each time point. Adding SWG does not affect the arrival of CD59 at the Golgi, but strongly delays its exit. This is particularly visible in the temporal projection (**Fig 1H**, right panel) showing that CD59 accumulates massively in the perinuclear region over time. Next, we performed RUSH assay with cells treated with siRNA against OSBP or with control siRNA (**Fig S1A**). Interestingly, the silencing of OSBP could partially recapitulate the effect observed with SWG: it does not modify the entry of CD59 at the Golgi, but delays its exit (**Fig S1B**). Altogether, these experiments suggest that SWG affects the formation of post-Golgi (but not pre-Golgi) transport intermediates, probably by targeting directly OSBP. Indeed, OSBP transfers cholesterol from the ER to the TGN via membrane contact sites while bypassing the cis Golgi (22), it is therefore in the best position to mediate the effects of SWG.

### SWG inhibits OSBP lipid exchange activity

To directly assess the effect of SWG on the lipid transfer activity of OSBP, we conducted in vitro liposome-based reconstitution experiments. We first monitored the transfer of a cholesterol fluorescent analogue (dehydroergosterol, DHE) from liposomes mimicking the ER (L_E_) to liposomes mimicking the Golgi (L_G_) (**Fig 2A**). To facilitate the analysis, we used a short OSBP construct (residues 401-807), corresponding to the ORD domain, which transfers rapidly DHE between liposomes (**Fig 2B**, grey trace). We observed that this rate decreased drastically as we increased the concentration of SWG in the reaction mix (**Fig 2B**). SWG blocked ORD-catalyzed transfer of DHE with an apparent K_i_ of <0.1 nM (**Fig 2C**), suggesting that SWG binds to the ORD domain of OSBP with a very high affinity. Next, we measured the transfer of PI(4)P by the ORD from L_G_ to L_E_ liposomes using a specific probe (PH^FAPP^) labeled with a NBD molecule, as previously described (5) (**Fig 2D**). The fluorescence of the probe was high at the surface of L_G_ liposomes, but was subsequently quenched as the probe followed PI(4)P relocation to L_E_ liposomes doped with Rhodamine lipids (**Fig 2E**). As with sterol transfer, SWG decreased the PI(4)P transfer rate in a dose-dependent manner, indicating that this drug inhibits the two transfer steps of the OSBP cycle.

**Figure 2:**
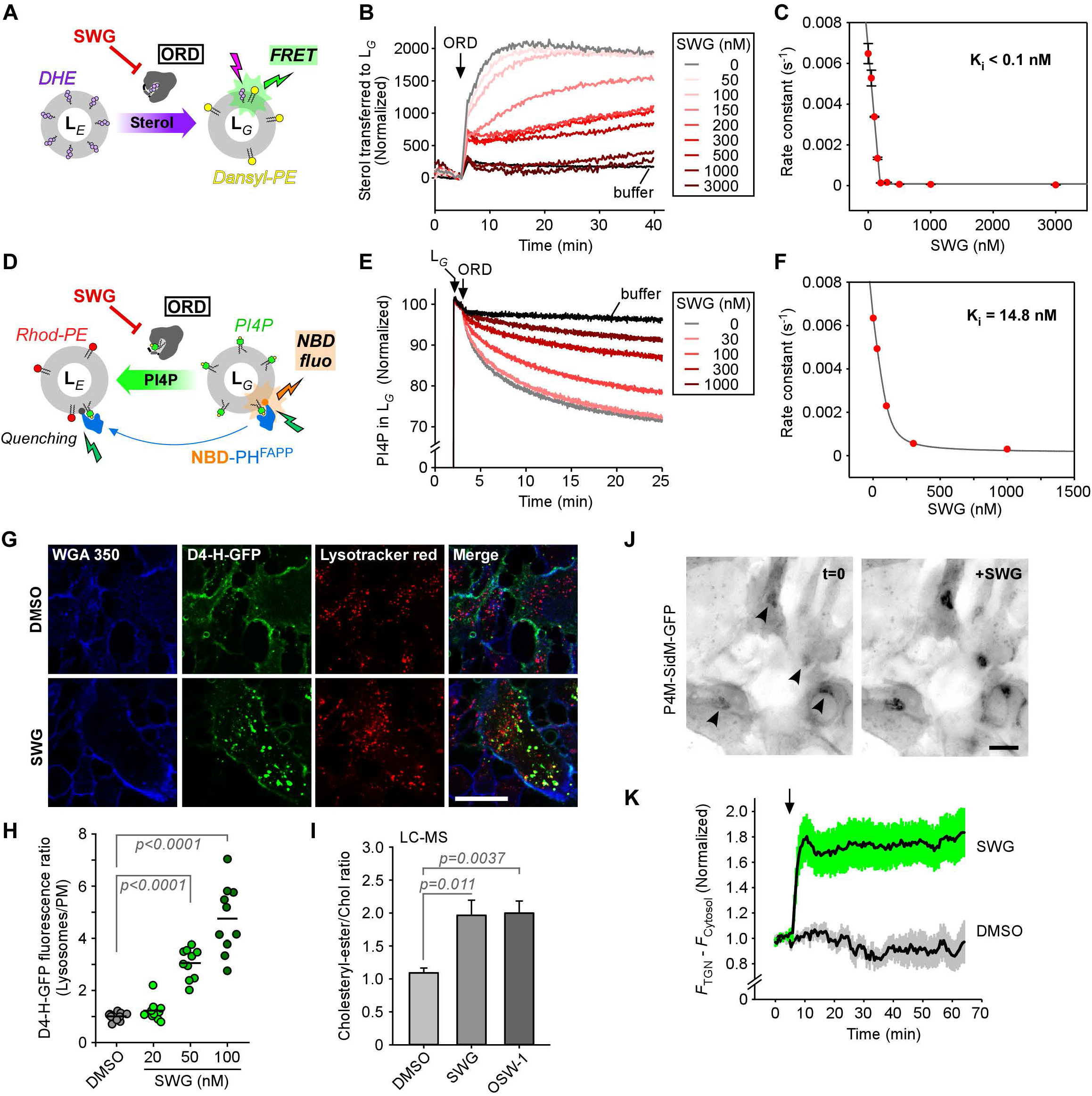
SWG is a potent inhibitor of the OSBP lipid exchange cycle and affects cellular lipid distribution. **A.** Experimental strategy for sterol transfer assay. **B.** DHE transfer assay between L_E_ (130μM) and L_G_ (130 μM) in the presence of OSBP ORD domain (0.2 μM) with increasing amounts of SWG. **C.** Inhibitory effect of SWG on DHE transfer mediated by OSBP ORD domain. The graph represents the rate constants obtained from fitting curves such as in B from three independent experiments, as a function of SWG concentration. The curve is fitted with a quadratic equation, which gives the *K*_i_ (see experimental procedures). **D.** PI(4)P transfer assay. **E.** PI(4)P transfer measurement between L_G_ (300 μM) and L_E_ (300μM) in the presence of NBD-PH^FAPP^ (0.3 μM) and OSBP ORD domain (0.1 μM) with increasing amounts of SWG. **F.** Inhibitory effect of SWG on PI(4)P transfer. **G.** Confocal images of RPE-1 stably expressing D4-H-GFP and either treated with SWG (100 nM) or DMSO for 16 h. Thereafter, the cells were fixed and labelled with WGA-Alexafluor 350 (plasma membrane marker) and Lysotracker red (lysosomal marker). **H.** D4-H-GFP fluorescence intensity ratio (lysosome/PM) from n = 10 measurements for each concentration condition. Significance tested with an unpaired t-test. **I.** Cholesteryl-ester/Cholesterol ratio measured by LC-MS from lipid extracts of RPE-1 treated with SWG (50 nM) or OSW-1 (20 nM) for 2 h. Results are means ± SEM (n=3). **J.** Time-lapse microscopy of RPE-1 stably expressing P4M-SidM-GFP and treated with SWG (50 nM) as indicated. Real-time images were acquired using a wide-field microscope at 2 frames/min. **K.** Time course of P4M-SidM-GFP at the TGN (the specific P4M-SidM-GFP level at the TGN shown is obtained by subtracting the cytosol value from the TGN value). The arrow indicates SWG addition into the cell medium. Data are means ± SEM (n=6).

We have previously shown that OSBP contributes significantly to the intracellular sterol distribution and influences the lipid order gradient between membranes along the secretory pathway (6). For this purpose, OSBP consumes massive amounts of PI(4)P at the TGN. We therefore tested whether SWG affects cholesterol distribution and PI(4)P turnover in cells. To do this, we first used RPE-1 cells stably expressing the probe D4-H-GFP derived from the Perfringolysin O theta toxin of *Cl. perfringens*, which specifically recognizes cholesterol from 20 mol% in membranes. Previous work has shown that D4-H decorates mainly the plasma membrane at steady state, but that following cholesterol depletion (e. g. caused by the extracellular addition of methyl-β-cyclodextrin), it translocates to endosomal compartments (23). Accordingly, we found that D4-H-GFP was mostly at the plasma membrane (positive for WGA) in control cells (**Fig 2G**). Then, upon treatment with SWG, D4-H-GFP redistributed to endo-lysosomes in a dose-dependent manner, indicating that cholesterol levels dropped at the plasma membrane (**Fig 2H**). To further examine whether SWG affects other pools of cholesterol, we measured the cholesteryl-ester/cholesterol ratio by LC/MS. Indeed, we suspected that, by inhibiting OSBP, SWG could lead to excessive amounts of cholesterol in the ER, which would foster its esterification and incorporation into lipid droplets. As shown in **Fig 2I**, a two-hour treatment with 100 nM of SWG significantly increased the cholesteryl-ester/cholesterol ratio, by about 2-fold, as compared to control DMSO-treated cells (**Fig 2I**). As a comparison, we also tested OSW-1, another OSBP inhibitor, which provided a similar effect. Therefore, these two drugs seem to promote cholesterol esterification, which is consistent with a blockage of OSBP activity. Because OSBP continuously consumes PI(4)P at the TGN, its pharmacological inhibition should protect a substantial fraction of PI(4)P, as previously reported (6). To verify this, we monitored the distribution of the PI(4)P probe P4M-SidM coupled to a fluorophore by live cell imaging. P4M-SidM-GFP stably expressed in RPE-1 cells was mostly cytosolic and decorated slightly the TGN, in control cells (**Fig 2J**). However, the addition of SWG caused a sudden twofold increase in TGN-associated P4M-SidM-GFP fluorescence, suggesting that the drug is capable of rapidly crossing cell membranes and acting on the PI(4)P pool of the TGN (**Fig 2J, K**). Altogether, these experiments suggest that SWG affects membrane cholesterol levels at multiple steps along the secretory pathway, which coincides with an increase in the level of PI(4)P at the Golgi, suggesting that OSBP is the key cellular target of SWG.

By controlling the turnover of PI(4)P at the Golgi, OSBP regulates its own membrane attachment and ER-TGN contact formation (5). It is therefore possible that SWG facilitates organelle tethering by inhibiting OSBP. To verify this, we used cells overexpressing mCherry-OSBP, GFP-VAP-A and a TGN marker, TagBFP-βGalT1. We observed that following the addition of SWG, not only OSBP, but also VAP-A, translocate rapidly and massively to the perinuclear region, suggesting that SWG promotes the formation of ER-TGN contact sites (**Fig S2**). Note that this observation corroborates our data showing that endogenous OSBP shifted to the TGN after a short treatment of SWG (**Fig 1F**). This effect is probably due to the inhibition of OSBP exchange activity, but it cannot be excluded that SWG also affects structural determinants of contact sites.

### Imaging SWG reveals its specificity for OSBP

We suspected that SWG could be intrinsically fluorescent because its stilbene motif comprises several conjugated double bonds (**Fig 1A**). To test this, we performed fluorimetric analysis of the drug in a saline buffer. As shown in **Fig 3A**, SWG exhibited excitation and emission spectra in the near UV range, with excitation and emission peaks at 330 and 410 nm, respectively. We then sought to visualize SWG in a cellular context *via* its intrinsic fluorescence. Using an imaging system adapted to UV, we directly assessed SWG subcellular distribution in RPE-1 cells after a treatment of 30 min at 37°C with 1 μM of the drug. Remarkably, SWG was enriched exactly where OSBP is located, that is, in the TGN region marked by TGN-46 (**Fig 3B**). Outside the TGN, SWG showed a weaker and diffused signal, as it likely labels membranes due to its partial hydrophobicity. Importantly, SWG distribution contrasted with that of a previously reported fluorescent SWG derivative (called SG-Fluor), consisting of SWG tagged with an additional fluorescent group, which decorated endosomal compartments (18). We next monitored the incorporation of SWG into the TGN over time (**Fig S3A, 3C**). Interestingly, after 5-10 minutes of incubation, SWG fluorescence was already visible in the perinuclear region, suggesting that SWG rapidly crosses the plasma membrane. This was confirmed by experiments showing that SWG could readily diffuse through giant plasma membrane vesicles (**Fig S3C, S3D**). The gradual increase (t_1/2_ = 25 min) of SWG at the TGN level coincided with the recruitment of OSBP in the same region over time (**Fig 3C, S3B**).

**Figure 3:**
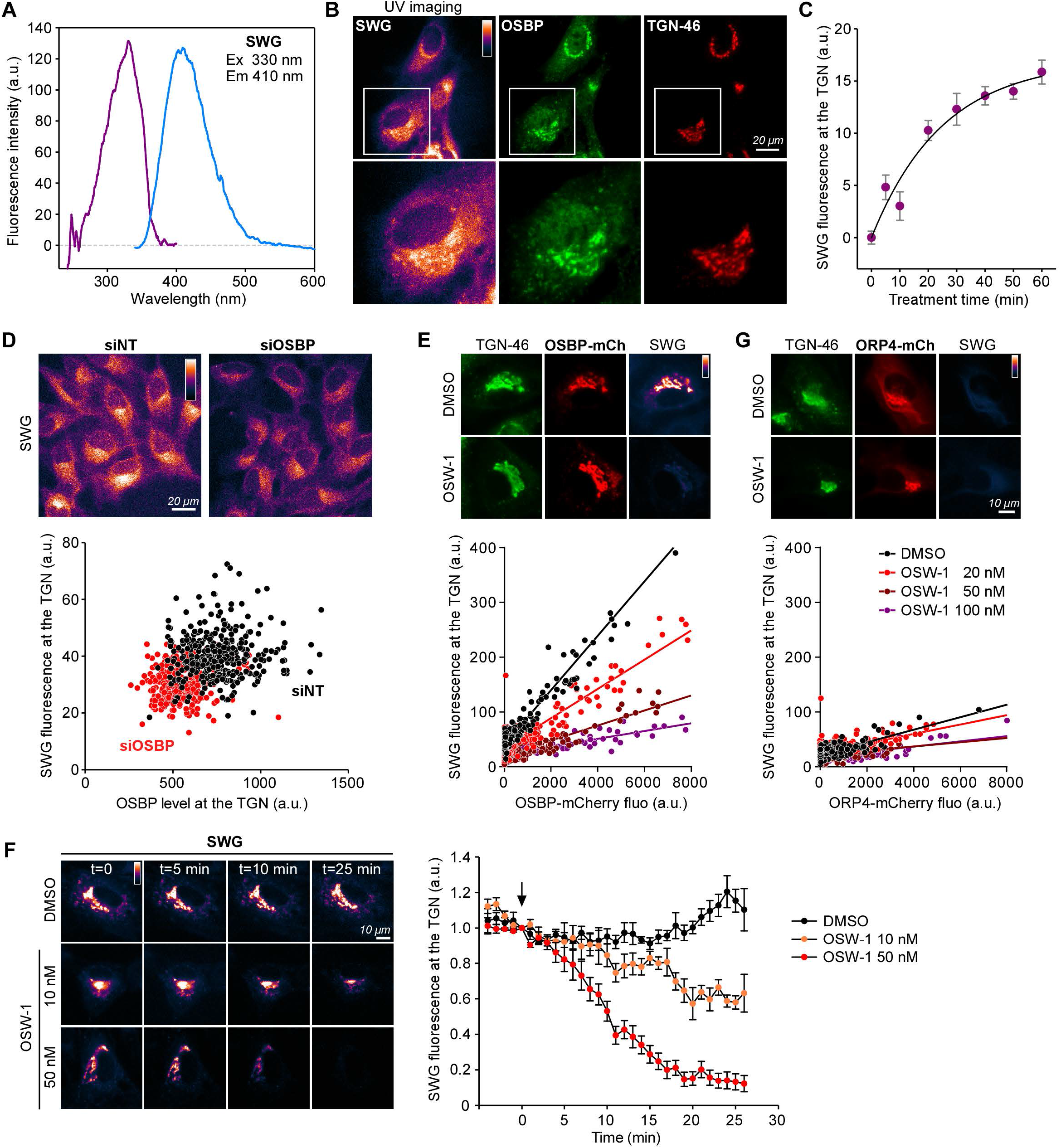
Direct fluorescent imaging of SWG in cells provides evidence of its specificity for OSBP. **A.** Excitation and emission spectra of SWG. **B.** Epifluorescence images of RPE-1 cells labelled with SWG (1 μM) for 30 min in growth medium at 37°C, fixed, permeabilized and processed for immunofluorescence to assess the localization of endogenous OSBP and TGN-46. Scale bar: 20 μm. **C.** SWG fluorescence intensity at the TGN over time. Data are means ± SEM (n=10). **D.** RPE-1 cells were treated either with control siRNA (siNT) or siRNA against OSBP, then processed as in B. Top: representative epifluorescence images of SWG. Bottom: SWG fluorescence at the TGN as a function of OSBP cellular levels, as determined by immunofluorescence quantification. Measurements were performed on more than 300 cells for each condition. **E.** Top: Epifluorescence images of RPE-1 transfected with OSBP-mCherry for 18 h, labelled with SWG (1 μM) for 30 min in the presence of OSW-1 (100 nM) or DMSO as control, then immunolabelled with anti-TGN-46. Bottom: SWG fluorescence intensity at the TGN as a function of OSBP-mCherry expression levels in the presence of indicated amount of OSW-1. Measurements were performed on 200–325 cells for each condition. **F.** RPE-1 cells were transfected with ORP4-mCherry for 18 h and treated as in E. **G.** Left: Time-lapse microscopy of RPE-1 cells overexpressing OSBP-mCherry, labelled with SWG (1 μM) for 30 min and treated with OSW-1 or DMSO as indicated. Right: Normalized levels of SWG at the TGN. The arrow indicates drug addition into the cell medium. Data are means ± SEM (n=3).

We reasoned that if SWG binds specifically to OSBP, the cellular SWG fluorescence intensity should reflect properly the abundance of OSBP. We therefore assessed SWG fluorescence in cells in which OSBP was either silenced or overexpressed (**Fig 3D, E**). We treated RPE-1 cells with siRNA directed against OSBP or with control siRNA for 72 h and then incubated the cells with 1 μM SWG for 30 min. Although silencing efficiency was variable among cells (**Fig 3D**, bottom panel), SWG fluorescence at the TGN was generally much less intense in silenced cells as compared to control cells, indicating that SWG is poorly recruited when low amounts of OSBP are present. Conversely, in OSBP overexpressing cells, SWG was extremely enriched at the TGN and co-localized with OSBP-mCherry and TGN-46 (**Fig 3E**). The level of SWG fluorescence in the cells almost perfectly correlated with that of OSBP-mCherry, suggesting that incorporation of SWG at the TGN is attributable to OSBP. To further verify this hypothesis, we treated the cells with SWG and an increasing dose of OSW-1. This compound also binds to OSBP but is not fluorescent, so it should compete with SWG for OSBP. OSW-1 prevented the TGN labelling by SWG in a dose-dependent manner. The addition of 20, 50 and 100 nM of OSW-1 caused a reduction in the amount of SWG at the TGN by 1.9, 3.6 and 7-fold, respectively (**Fig 3E**, bottom panel). Alternatively, we first labelled cells overexpressing OSBP-mCherry with SWG and then added the competitor OSW-1 (**Fig 3F**). By time lapse imaging experiment, we observed that OSW-1 gradually chased out SWG from the TGN in a dose-dependent manner (t_1/2_ = 9.2 ± 0.7 min with 50 nM OSW-1), further indicating that the retention of SWG at the TGN is most likely related only to OSBP.

ORPphilins were initially described as being able to bind to OSBP and ORP4, however, schweinfurthins A and B exhibited a clear preference for OSBP over ORP4 (1). In order to assess SWG specificity for OSBP over ORP4, we overexpressed ORP4-mCherry instead of OSBP-mCherry and performed treatment with SWG for imaging (**Fig 3G**). ORP4 did not facilitate SWG recruitment, thus demonstrating that SWG selectively targets OSBP.

### Biochemical characterization of SWG

Because SWG rapidly penetrates into cells to target OSBP (**Fig S3**), we hypothesized that SWG is able to passively interact with membranes due to its partial hydrophobicity. To characterize the interactions of SWG with both membranes and with OSBP, we performed in vitro fluorescence measurements of SWG. Spectral analysis revealed that when the drug was switched from an aqueous medium to polar solvents (such as methanol or acetonitrile), its fluorescence increased and this effect was accompanied by a blue shift (**Fig 4A**). Instead, when SWG was placed in a nonpolar solvent such as hexane, no fluorescence could be detected. Interestingly, adding detergent micelles (thesit; green curves) or liposomes (**Fig 4B**, blue curve) further increased the fluorescent signal and structured SWG emission spectrum, indicating that SWG fluorescence is even more sensitive to interfacial environments.

**Figure 4:**
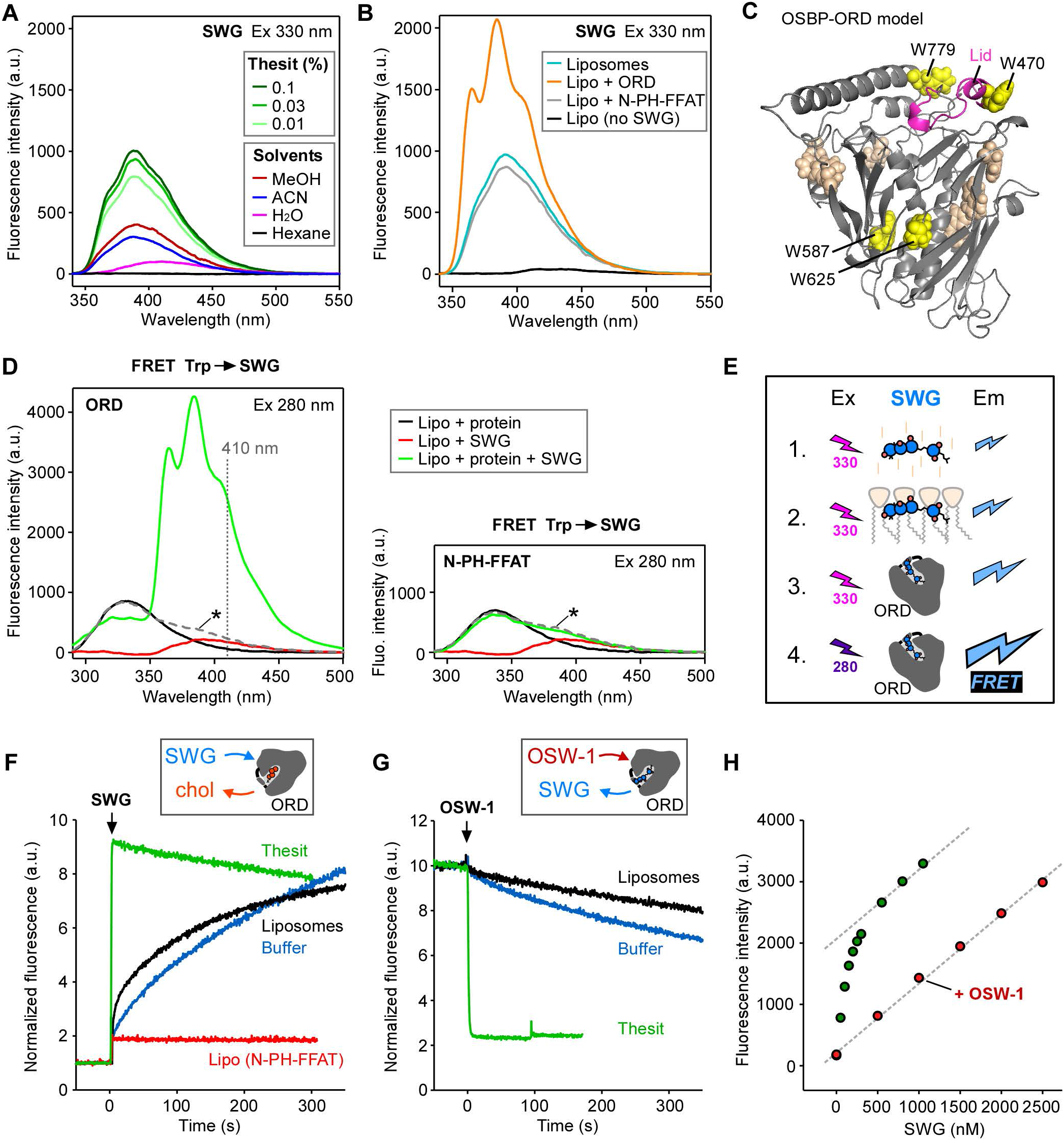
The fluorescence properties of SWG are influenced by its environment. **A.** Emission spectra of SWG (200 nM) upon excitation at 330 nm in various solvents or in the presence of the indicated amount detergent in water (green curves). **B.** Emission spectra of SWG (200 nM) in the presence of liposomes (0.1 mg/mL), with or without the addition of ORD (200 nM) or N-PH-FFAT (200 nM) in HKM buffer. The black curve represents the signal obtained with liposomes without SWG. **C.** Homology modelling of the ORD domain of OSBP. Tryptophan residues (represented as spheres) are colored in yellow when close to the lipid-binding cavity. The lid of the ORD is colored in pink. **D.** Spectra analysis upon tryptophan excitation at 280 nm. The protein added was either the ORD (200 nM, left panel) or the N-PH-FFAT construct (200 nM, right panel) in HKM buffer, in the presence of liposomes (0.1 mg/mL), with or without the addition of SWG (200 nM). The dashed curves (asterisk) represent the theoretical sum of the fluorescence values from black and red curves. **E.** Sketch summarizing the effects of the local environment on SWG fluorescence intensity. **F.** Time course of SWG (200 nM) binding into the ORD (200 nM) as measured by FRET (Ex 280 nm; Em 410 nm). When indicated, the reaction was performed in the presence of thesit (0.1 %, green curve), liposomes (0.1 mg/mL, black curve) or in buffer HKM alone (blue curve). In a control experiment, the ORD was replaced with N-PH-FFAT (200 nM, red curve). **G.** Time course of SWG (200 nM) release from the ORD (200 nM) using OSW-1 (500 nM). The conditions are the same as in F. **H.** Binding of SWG to ORD measured by FRET. The sample initially contained the ORD (200 nM) in thesit (0.1 %), to which increasing amount of SWG was added (green dots). When indicated, the initial sample was supplemented with 200 nM OSW-1 (red dots).

We reasoned that SWG could also experience a change in environment if it enters in the hydrophobic cavity of the ORD of OSBP. We added OSBP constructs to a mixture containing 200 nM of SWG and liposomes (0.1 mg/mL), and recorded SWG emission spectra upon excitation at 330 nm (**Fig 4B**). Remarkably, the addition of ORD (200 nM) more than doubled the fluorescence intensity of SWG as compared to liposomes alone. In contrast, adding 200 nM of the N-PH-FFAT construct, i.e., the N-terminal region of OSBP, did not promote any fluorescence change. These results strongly suggest that the drug inserts into the binding cavity of the ORD. To further characterize the contact between SWG and OSBP, we performed the 3D structure prediction of the ORD of OSBP by homology modelling, based on the crystal structure of the yeast OSBP-related protein Osh1 (24), which also exchanges sterol for PI(4)P (**Fig 4C**). We noticed that several tryptophan residues are close to the cavity of the ORD (indicated in yellow) and thus in good position to form fluorescence resonance energy transfer (FRET) pairs with SWG. To test this hypothesis, we mixed liposomes with the ORD, with SWG, or with a mixture of ORD and SWG, and performed spectral analysis upon tryptophan excitation at 280 nm (**Fig 4D**). As expected, a tryptophan fluorescence signal peaking at ~340 nm was observed when the ORD was employed alone (black line), whereas the signal obtained with SWG alone (red line) was weak since it is poorly excited at 280 nm (**Fig 3A**). Strikingly, when both the ORD and SWG were present, we observed a massive FRET signal (green line) with peaks characteristic of SWG (as seen in **Fig 4B**). The FRET signal was much higher than the theoretical sum of the signals obtained with the ORD or SWG separately (dashed grey line). The FRET signal was also 2-fold greater than the SWG fluorescent signal obtained upon direct excitation at 330 nm under the same conditions. The apparition of FRET was concomitant with partial quenching of tryptophan emission at 340 nm. Altogether, these results provide evidence that SWG interacts with the ORD of OSBP by entering in its lipid binding pocket, and also suggest that FRET between the ORD and SWG is a useful tool to probe this interaction (**Fig 4E**).

### Kinetics of SWG binding to the ORD

We took advantage of the large FRET signal to monitor the binding and dissociation kinetics of SWG to the ORD. We first mixed SWG with ORD (200 nM each) and measured the changes in fluorescence over time at 410 nm, which is the most specific FRET wavelength, upon tryptophan excitation at 280 nm. As shown in **Fig 4F**, SWG binding to the ORD was faster when liposomes (1 mg/mL) were present in the cuvette than simply in buffer alone (compare the black and blue curves). As a control, we replaced the ORD by the N-terminal half of OSBP (residues 1-408, N-PH-FFAT) and observed no change in fluorescence (red curve). Interestingly, addition of detergent micelles (thesit) caused the reaction to be completed in about 2 seconds (green curve, see also **Fig S4A**), suggesting that the association rate of SWG with the ORD is very fast (k_on_ > 2 s^−1^) in the presence of an interfacial environment. The effect of membranes or micelles can be explained by the fact that the ORD is purified with a cholesterol molecule in its binding pocket, as shown by mass spectrometric analysis (**Fig S4B**). The presence of a membrane interface likely facilitates the dissociation of cholesterol from the ORD, thereby allowing SWG entry. Our results in **Fig 3F** suggest that SWG could be removed by OSW-1 from their binding site, so we next monitored the effect of OSW-1 addition (in slight excess; 500 nM) on SWG-loaded ORD in the same conditions as in **Fig 4F**, i.e., in buffer supplemented or not with liposomes or micelles. With micelles, the rate of dissociation of SWG from the ORD was very fast (k_off_ > 2 s^−1^), in contrast, SWG exit from the ORD in buffer alone or in the presence of liposomes was rather slow (**Fig 4G**). Finally, to further analyze the interaction of SWG with the ORD of OSBP, we performed equilibrium measurements using FRET. **Fig 4H** shows a titration experiment in which we added increasing amounts of SWG to a mixture containing the ORD (200 nM) and 0.1% thesit. Below 200 nM, the stepwise addition of SWG induced a large FRET signal. Thereafter, the apparent increase in fluorescence was no longer due to FRET but to SWG direct fluorescence. To reveal the saturation of the FRET signal, we repeated the same experiment, but in the presence of OSW-1 (200 nM) to block the ORD. In this case, the increase in fluorescence came only from SWG increments in the mixture and not from FRET. Fitting the dose-response curve after correction of the non-specific signal indicated an apparent affinity of SWG for the ORD in the presence of detergent micelles of K_d_ = 35.5 nM. This affinity is much lower than the K_i_ observed in cholesterol exchange assays performed with liposomes (**Fig 2C**) suggesting that the micellar environment unlocks the ORD, hence leading to a decrease in affinity and to a dramatic acceleration in the dissociation kinetics of ORD ligands.

## Discussion

The discovery of ORPphilins is an important step for the study of ORPs, highlighting the importance of these proteins in cancer, and providing pharmacological tools to characterize their molecular activities and physiological functions. ORPphilins include compounds with a variety of structures, including OSW-1, cephalostatin, stelletin and schweinfurthin sub-families (1). Although they are described to have the same intracellular targets (OSBP and ORP4), their differential affinity and the way they interact with the cellular environment could have different functional and physiological consequences.

The present work addresses several properties of SWG, a natural product of the schweinfurthin family, which displays antiproliferative activities for specific tumor-derived cell lines. By using a combination of reconstitutions on artificial membranes and cellular approaches, we show that SWG binds directly to the ORD domain of OSBP. This binding prevents the lipid exchange cycle of OSBP, i.e., PI(4)P consumption and cholesterol transfer, and promotes ER-TGN contact site stabilization. As a result, post-Golgi trafficking and plasma membrane cholesterol levels are severely affected.

We showed that when there are more or less copies of OSBP present in a cell type, a higher or lower SWG concentration is required, respectively, to mediate a cytotoxic effect (**Fig 1C**). This probably reflects a buffering effect: SWG concentration must be adjusted to reduce total OSBP activity below a critical threshold for cell viability. Consistently, it is reported that OSBP overexpression confers cell resistance to schweinfurthins, while OSBP silencing sensitizes cells to these drugs (1). First, this excludes a potential toxic effect due to excessive ER-TGN membrane tethering triggered by SWG, because in this case, cytotoxicity would increase with overexpression of OSBP. Second, this may explain the specific cytotoxicity of these drugs for a subset a cancer cell lines. In the future, it would be interesting to verify whether cell sensitivity to various ORPphilins could be predicted according to tissue concentration of OSBP.

High levels of cholesterol in the ER usually leads to two cellular responses: the rapid esterification of excess cholesterol followed by its incorporation into lipid droplets (25), and a slower transcriptional response mediated by the cholesterol homeostatic regulatory machinery (26), including the transcription factors sterol regulatory element-binding protein (SREBP) and liver X-receptor (LXR), which stop cholesterol biosynthesis and uptake, and promote its export from the cell. The addition of SWG blocks cholesterol fluxes through OSBP coming from the ER. As a result, we showed an increased cholesterol esterification. Interestingly, previous work has revealed that treatment with 3-deoxyschweinfurthin B induces changes in the expression of genes related to cholesterol homeostasis controlled by SREBP and LXR (17). Because these regulators are coupled to cholesterol-sensing mechanisms localized at the ER, it is possible that inhibiting OSBP triggers such homeostatic response. Further studies would be needed, however, to address the direct involvement of the OSBP cycle in cholesterol sensing at the ER (e.g. through the SCAP/SREBP-2 pathway).

Fluorescent probes are instrumental for tracking biomolecules with good spatio-temporal resolution in living cells by imaging. In particular, the use of fluorescently tagged drugs can shed light on their specific interactions with cellular receptors, and provide molecular insights into pharmacological mechanisms. The major drawback of adding a large fluorescent moiety to compounds is that it can alter their pharmacological properties by modifying their chemical properties and/or their cellular distribution. Several studies have been conducted to make some ORPphilins fluorescent in order to visualize them in cells. A fluorescent OSW-1 analogue, labelled with a 4-(N,N-dimethylaminosulfonyl)-7-N-methylamino)-2,1,3-benzoxadiazolyl (DBD) group, was shown to distribute at the Golgi and to a lesser extent at the ER, and retains anti-proliferative properties (27). For schweinfurthins, fluorescent probes with differential activity between cell lines have been synthesized, such as schweinfurthin B and F modified by a *p*-nitro stilbene group, showing fluorescence in the visible spectrum (19). However, these analogues localized to punctate structures that do not resemble the Golgi compartment. Similarly, a fluorescent probe consisting of SWG with an additional *p*-nitro stilbene group (called SG-Fluor) was localized to punctate structures that overlapped with the endosomal marker LAMP1 (18). This seems not consistent with the subcellular distribution of OSBP, which is mainly found at the TGN and weakly localized to endosomes. In the present study, we exploited the intrinsic fluorescence properties of SWG in imaging and in membrane reconstitution assays. By using a UV-sensitive imaging system, we were able to observe directly SWG in its intracellular site of action, that is, at the TGN. We showed that when OSBP cellular abundance was adjusted (via silencing or overexpression), the cellular incorporation of SWG was modified accordingly, suggesting a high specificity of the drug for OSBP. We also showed that variations in ORP4 levels had little or no impact on SWG cell labelling. These results indicate that SWG is selective towards OSBP. This is consistent with earlier findings showing that Schweinfurthin A had a better binding specificity to OSBP as compared to ORP4 (1).

The ORD domain of several ORP/Osh proteins, including OSBP, comprises tryptophans close to the lipid-binding pocket (**Fig 4C**). In a previous study, we showed that FRET could be measured between tryptophans of Osh4 and the intrinsically fluorescent sterol DHE once bound in the pocket (28). Here, we took advantage of the fluorescence properties of SWG, which absorption-emission spectra partly overlap with that of DHE, to perform equilibrium and kinetics measurements by using FRET between tryptophans of OSBP ORD and SWG. We have shown that the association and dissociation rates of SWG from ORD domain were very fast (>2 s^−1^). More specifically, SWG entry in the ORD was facilitated when membranes (or micelles) are present, which is probably due to the fact that the ORD is co-purified with a cholesterol molecule bound in the pocket. Our hypothesis is that the ORD must first interact with the membrane to allow cholesterol exit so that SWG can then enter the pocket. Understanding this mechanism will, however, require further investigation.

Finally, this work invites a comparison between the OSBP-targeting compounds SWG and OSW-1. Both drugs strongly inhibit OSBP-catalyzed lipid transfer with K_i_<1 nM and promote ER-TGN contact sites formation (6). We have shown that OSW-1 prevents TGN labelling by SWG when both drugs are added simultaneously to the cells, and that SWG can be removed from the TGN when OSW-1 is added afterwards, even at low doses. Last, OSW-1 could displace SWG from purified OSBP ORD in vitro. These results suggest that OSW-1 has a stronger affinity for the ORD than SWG, and/or OSW-1 dissociates more slowly from the ORD. In this regard, we reported an apparent affinity of SWG for the ORD in the presence of detergent micelles of K_d_~35 nM, which is rather low compared to the measured K_i_. Akin to what has been observed recently on Osh6 (29), the ORD lid is probably unlocked by the interfacial micellar environment, hence leading to faster exchange kinetics and lower affinities. Further studies of the kinetics of SWG binding to or dissociation from the ORD with liposomes of defined composition should give information on the interfacial interactions involved in lipid exchange. The lack of fluorescence of OSW-1 and its too high affinity for the ORD, makes this inhibitor an almost irreversible ligand and therefore less suited for mechanistic investigations than SWG.

Because most ORPphilins (including OSW-1) interact with both OSBP and ORP4, it has been suggested that their anti-proliferative effect is due to ORP4 targeting and not OSBP (7). Indeed, high expression of ORP4 in immortalized cell lines, and in leukemia, leads to cell proliferation (30, 31). Moreover, a ~90% reduction in OSBP level does not alter cell proliferation (1). Intriguingly, our data suggest that SWG, which is a potent inhibitor of cell proliferation, selectively targets OSBP and not ORP4. In addition, our study provides a correlation between the level of OSBP expression and the differential sensitivity to SWG, suggesting that OSBP is a potential mediator of cell proliferation. Interestingly, a recent study has shown the direct activation of mTORC1 by OSBP-mediated cholesterol transfer, thereby clearly defining a link between OSBP activity and cell growth signaling (32). SWG might provide a useful tool to dissect this pathway.

## Experimental procedures

### Plant material and SWG isolation

The green fruits of *M. tanarius* were collected in June 2014 at A Lưới, Thừa Thiên-Huế Province, Vietnam, and authenticated by N.T. Cuong and D.D. Cuong. A voucher specimen (VN-2371) has been deposited at the Herbarium of the Institute of Ecology and Biological Resources of the Vietnam Academy of Science and Technology, Hanoi, Vietnam. Isolation of SWG was performed as previously described (33).

### Cell culture

hTERT-RPE1 cells (ATCC) were cultured in DMEM/F12 medium with glutaMAX (Gibco) containing 10% fetal calf serum, 1% antibiotics (Zell Shield, Minerva Biolabs) and were incubated at 37°C in a 5% CO2 humidified atmosphere. RPE1 cells stably expressing EGFP-D4H were selected using G418 (Sigma). Surviving colonies were isolated using cloning cylinders (Bel-Art), expanded, and further sorted by FACS (FACSAria III, BD Biosciences). RPE1 cells stably expressing EGFP-D4H or EGFP-P4M-SidM (6) were cultured in medium supplemented with G418 (500 μg/ml). Cells stably expressing Streptavidin-KDEL (hook) and SBP-EGFP-CD59 (reporter) (gift from G Boncompain, S Miserey-Lenkei and F Perez) were used for RUSH assays. Secretion was triggered by the addition of 80 μM biotin. For microscopy, cells were seeded at suitable density to reach 50–90% confluence on the day of imaging.

### Cell viability assay

Cells were seeded at 3000 cells/well density into white wall 96-well plates then treated on the following day with either SWG or Doxorubicin. After 72 h treatment, cell viability was measured using CellTiter-Glo Luminescent Cell Viability Assay (Promega) and the luminescence was detected in a FLUOstar Omega microplate reader (BMG Labtech). Viability values of DMSO treated cells were considered as 100%, then surviving curves and IC50 values were determined using GraphPad Prism 7.0 software.

### Protein expression and purification

The human OSBP ORD fragment (401-807) with a C-terminal 6×His tag was purified from baculovirus-infected Sf9 cells. For cloning and protein expression, see Supplementary experimental procedures. Cell pellets were resuspended in lysis buffer (20 mM Tris pH 7.5, 300 mM NaCl, 20 mM imidazole, EDTA-free protease inhibitors and phosphatases inhibitors) and lysed with a Dounce homogenizer. After ultracentrifugation (125 000 g), OSBP-ORD from the supernatant was adsorbed on an HisPur™ Cobalt Resin (Thermo Scientific), submitted to 3 washes with lysis buffer supplemented with 800, 550, and 300 mM NaCl, respectively, and then eluted with 250 mM imidazole-containing buffer. OSBP-ORD fractions were pooled, concentrated on Amicon Ultra (cut-off 30 kDa), then purified on a Sephacryl S300 HK16/70 column (GE Healthcare) using an AKTÄ chromatography system (GE Healthcare). All steps were performed at 4°C. The purified protein fractions were pooled, concentrated, supplemented with 10% glycerol, flash-frozen in liquid nitrogen and stored at −80°C. The preparations of NBD-PH^FAPP^ and OSBP N-PH-FFAT (1-408 fragment) have been described previously (5, 34).

### Sterol transfer assay

The lipid compositions of the ER and Golgi liposomes were PC, PE, PS, PI, Dansyl-PE (64/19/5/12/10 mol %) and PC, PE, PS, PI, DHE (61.5/19/5/12/2.5 mol %). Measurements in triplicate were carried out in a TECAN Infinite M1000 Pro Microplate Reader using 96-well microplates (200 μL per well) equilibrated at 37°C, with the following settings: Ex 310/5 nm, Em 525/5 nm. Each well initially contained Golgi liposomes (130 μM lipid), ORD (0.2 μM) and SWG (as indicated) in 50 mM HEPES pH 7.2, 120 mM potassium acetate, 1 mM MgCl_2_ (HKM buffer). ER liposomes (130 μM lipid) were then added at the indicated time. Maximal exchange was controlled with 1mM methyl-β-cyclodextrin. Data analysis: The signal was first corrected from SWG intrinsic fluorescence, then curves were fitted either with a linear equation (for slow kinetics when SWG is >200 nM) or an exponential equation to determine the apparent kinetic constant, *k*. Because SWG has a strong affinity for ORD, the concentration of free SWG cannot be approximated to the total concentration of SWG. Therefore, the inhibition curve was fitted with a quadratic equation to obtain *K*_i_:

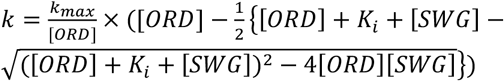

### PI(4)P transfer assay

The lipid composition of the ER and Golgi liposomes were PC, PE, PS, PI, rhodamine-PE, cholesterol (64/19/5/10/2/15 mol%) and PC, PE, PS, PI, PI(4)P(64/19/5/10/2mol%). Measurements were carried out in a Jasco FP-8300 spectrofluorimeter using a cylindrical quartz cuvette (600 μL) equilibrated at 37°C and equipped with a magnetic bar for continuous stirring. The cuvette initially contained NBD-PH^FAPP^ (300 nM) in HKM buffer. NBD emission was measured at 510 nm (excitation 460 nm). Golgi liposomes (300 μM lipid), ER liposomes (300 μM lipid) and ORD (0.1 μM) were then sequentially added at indicated times.

### Microscopy

Widefield microscopy was performed using an Olympus IX83 inverted microscope equipped with a Z-drift compensator, a scanning stage SCAN IM (Märzhäuser), and an iXon3 camera (Andor). For time-lapse imaging, cells plated in μ-Dish35 mm (Ibidi) were mounted in a stage chamber set at 37°C (Okolab). TagBFP, EGFP, and mCherry were observed using Chroma fluorescence filter sets (ref. 49000, 39002, 39010). Multidimensional acquisition and analysis was performed with MetaMorph software (Molecular Devices). Cells were placed in an appropriate initial volume of medium (phenol red-free, HEPES; Gibco) so that homogenous mixing of compounds (freshly diluted before use at 2–5 × concentration) was achieved upon subsequent additions during the time course measurements. Confocal microscopy was performed with a Zeiss LSM 780 microscope operated with ZEN software using a Plan-Apochromat 63X/1.4 Oil objective (Carl Zeiss).

### SWG fluorescence imaging

RPE-1 cells labelled with SWG were placed in phenol red-free DMEM/F12 with HEPES (Gibco) and observed using an IX83 inverted microscope (Olympus) equipped with an iXon3 blue-optimized EMCCD camera (Andor) and an UPlanSApo 60X/1.35 Oil objective (Olympus). SWG was imaged using Semrock BrightLine filters (320/40 nm band-pass filter, 347 nm dichroic beam splitter, and 390/40 nm band-pass filter).

### SWG fluorescence spectroscopy

SWG excitation (Ex 230-400/1 nm; Em 410/5 nm), SWG emission (Ex 330/1 nm; Em 340-600/5 nm) or FRET (Ex 280/1 nm; Em 340-600/5 nm) spectra were carried out in a Jasco FP-8300 spectrofluorimeter using a cylindrical quartz cuvette (600 μl) equilibrated at 37°C and equipped with a magnetic bar for continuous stirring. SWG (200 nM) emission spectra were measured in diverse environments as indicated. When indicated, ORD was added at 200 nM and liposomes (100% Egg PC) at 0.1 mg/ml. Kinetics FRET measurements were performed with the following settings: Ex 280/1 nm; Em 410/5 nm.

## Acknowledgements

We thank S. Abélanet, F. Brau, J. Cazareth and N. Leroudier (IPMC, Valbonne) for technical assistance; D. Lévy (Institut Curie, Paris), T. Virolle (Institut de Biologie Valrose, Nice) and M. Gutmann (SATT, Paris Saclay) for fruitful discussions; S. Miserey-Lenkei, G. Boncompain and F. Perez (Institut Curie, Paris) for sharing the RUSH cell lines.

## FOOTNOTES

This work was supported by the CNRS, Inserm, the Agence Nationale de la Recherche (ANR-15-CE11-0027-02 and ANR-11-LABX-0028-01), SATT Paris Saclay (Projet Glioved) and the International Associated Laboratory (LIA NATPROCHEMLAB) between the CNRS (ICSN, France) and the VAST (Institute of Marine Biochemistry, Vietnam). The Antonny lab is supported by a grant from the Fondation pour la Recherche Médicale (Convention DEQ20180339156 Equipes FRM 2018). M. Subra is supported by a Ph.D fellowship from the Université Côte d’Azur.

The abbreviations used are:

ER: endoplasmic reticulum
TGN: trans-Golgi network
LTP: lipid-transfer protein
OSBP: Oxysterol-binding protein
ORP: OSBP-related protein
ORD: OSBP-related domain
SWG: Schweinfurthin G
PI(4)P: phosphatidylinositol-4-phosphate
DHE: dehydroergosterol

