## Supporting Information for "Molecular and cellular dissection of the OSBP cycle through a fluorescent inhibitor"

This file includes:

- -Supplementary experimental procedures
- -Supplementary references
- -Figures S1 to S4

### Supplementary experimental procedures

#### *Reagents*

OSW-1 was a gift from Matthew D Shair (Harvard). Doxorubicin, methyl-β-cyclodextrin and G418 were purchased from Sigma-Aldrich. Lysotracker red (DND-99), Wheat germ agglutinin (WGA) Alexa Fluor conjugates were from Invitrogen.

#### *Antibodies*

Rabbit polyclonal antibody against OSBP (HPA039227) was from Sigma-Aldrich. Sheep polyclonal antibody against TGN-46 was from AbD Serotec (Bio-Rad). Antibody against β-Actin was from Sigma-Aldrich. Secondary Alexa Fluor-conjugated antibodies were from Invitrogen and secondary peroxidase-conjugated antibodies were from Jackson ImmunoResearch.

#### *Lipids*

Egg PC, brain PS, brain PI(4)P, liver PI, liver PE, dansyl-PE (1,2-dioleoyl-sn-glycero-3-phosphoethanolamine-N-(5-dimethylamino-1-naphthalenesulfonyl)) and rhodamine-PE [1,2-dipalmitoyl-sn-glycero-3-phosphoethanolamine-N-(lissamine rhodamine B sulfonyl)] were obtained from Avanti Polar Lipids. Cholesterol and dehydroergosterol were from Sigma-Aldrich.

#### *Liposomes*

Lipids in chloroform, or in chloroform:methanol (2:1) in the case of mixtures containing PI(4)P, were mixed at the desired molar ratio and the solvent was removed in a rotary evaporator. The lipid films were hydrated in 50 mM HEPES pH 7.2 and 120 mM potassium acetate (HK buffer, which was degassed before use) to give a suspension of large multilamellar liposomes (lipid concentration: 2-5 mM). The suspension was then frozen in liquid nitrogen and thawed in a warm water bath five times. Liposomes were extruded through 0.1 µm pore size polycarbonate filters using hand extruder (Avanti Polar Lipids) and were used within 1-2 days.

#### *Cell culture*

A549, MCF-7, PC3 (ATCC) and MDA MB 231 (DSMZ) cells were cultured in RPMI medium. U87, HeLa and MRC-5 cells (ATCC) were cultured in DMEM. All cell lines growth media were supplemented with 10% fetal calf serum. Cells were incubated at 37°C in a 5% CO2 humidified atmosphere.

#### *Plasmids and cell transfection*

The constructs pmCherry-N1 OSBP, pmCherry-N1 ORP4L, pEGFP-C1 VAP-A, pTagBFP-βGalT1 have been described previously (1–3). The domain 4 (D4) of the Perfringolysin O theta toxin of *Cl. perfringens* was PCR-amplified from the plasmid pET28-D4 (shared by Mitsuhiro Abe and Toshihide Kobayashi; RIKEN, Japan) and subcloned into pEGFP-C1. The D4-H, i.e., having the point-mutation D434S, was produced by Quickchange site-directed mutagenesis (Agilent). For protein expression, cells were transfected by electroporation using the Amaxa Nucleofector device (Lonza), for 18 to 24 h. For OSBP silencing, RPE-1 cells were transfected by electroporation (Amaxa, Lonza) with ON-TARGETplus Human OSBP siRNAs from Dharmacon (GE Healthcare) for 72 h, as previously described (2, 3). The OSBP target sequence is GCAAUGACUUGAUAGCUAA (Dharmacon). The non-targeting siRNA (si-NT) has the following sequence: UGGUUUACAUGUUGUGUGA (Dharmacon).

#### *Immunofluorescence*

Cells cultured on coverslips or on µ-slides (Ibidi) were washed once with PBS then fixed with paraformaldehyde (3%) and glutaraldehyde (0.1%) in PBS for 10 min. Fixed cells were soaked in PBS supplemented with $NH_4Cl$ (50 mM) for 10 min, and then permeabilized in permeabilization buffer (Saponin 0.05 %, BSA 0.2 %, PBS) during 30 min at RT°. Cells were then immuno-labeled with primary antibodies (diluted in permeabilization buffer) for 1 h at RT°, washed 3 times with permeabilization buffer, further incubated with secondary Alexa Fluor-conjugated antibodies (diluted in permeabilization buffer) for 30 min at RT°, rinsed and eventually embedded in Mowiol solution.

***Giant plasma membrane vesicles (GPMVs) preparation and analysis***

RPE1 cells were seeded one day before the experiment and grew to 70 % confluence. Cells were washed twice with GPMV buffer (10 mM HEPES pH 7.4, 150 mM NaCl, 2 mM $CaCl_2$) and incubated with GPMV buffer containing 25 mM PFA and 2 mM DTT at 37°C for 3 h (which was defined as the shortest timing to obtain enough GPMVs from RPE1). Then, the supernatant containing GPMV was collected. For microscopy, 8-well µ-slides (Ibidi) were coated with 10 µg/mL fibronectin overnight at 4°C with agitation. Then, 200 µL of a solution containing GPMVs labelled with 1 µg/mL WGA-Alexa 488 were placed on 8-well µ-slides to sit and stabilize at the bottom of the slide for 1 hour. Thereafter, 10 µL of SWG (5 µM final) were added to the GPMVs solution and the kinetics were recorded at 3 frames/min during 10 minutes with a widefield IX83 inverted microscope (Olympus).

***Image analysis***

Automated custom macros were written in Fiji (ImageJ, NIH) to perform the analysis whenever applicable. For SWG quantification, images were first background corrected, then TGN-46 images were used to generate binary masks by intensity thresholding. The in-built ‘analyze particles’ command was then used to measure the area and integrated pixel intensity of regions of interest in the corresponding SWG and OSBP images. For time-lapse imaging of SWG, a photobleaching correction factor was applied to each image. The photobleaching decay was calculated from regions of interest of images acquired with the same illumination settings as for the experimental conditions, but in this case without any addition. Photobleaching correction was done on each experimental day. For D4-H-GFP quantification, images were background corrected, then a binary mask defined by WGA-Alexa 350 was used to quantify the mean D4-H-GFP fluorescence intensity at the plasma membrane, and a binary mask defined by Lysotracker red was used to quantify the mean D4-H-GFP fluorescence intensity at lysosomes. The ratios of the means were then plotted. RUSH assay quantification was performed as described previously (4), except that the values were normalized to the initial value. GFP-P4M-SidM quantification was performed as described previously (2). Colocalization analyses were performed on 16-bit confocal images using the Volocity software (PerkinElmer). From single optical sections, off-cell regions were used for setting threshold values, then regions encompassing cells were manually traced and Pearson’s correlation coefficients were calculated for each of them.

***OSBP-ORD fragment cloning and expression in Sf9 cells***

The ORD fragment (401-807) of human OSBP was purified from baculovirus-infected Sf9 cells. The ORD sequence was inserted into the pFastBac HTA modified vector, as described previously (3). DNA sequence was PCR amplified using the pENTR/D-(full-length OSBP-thrombin site) as matrix and cloned into the BamHI-digested pFastBac HTA modified vector using the GeneArt$^{TM}$ Seamless Cloning and Assembly Kit (Invitrogen). Recombinant vectors were then transformed into DH10 Bac *E.coli.* Recombinant bacmids were selected as described in Bac to Bac$^{R}$ Expression System user manual (Invitrogen) and used to produce recombinant Baculovirus. For protein expression, SF9 cells cultured at 27°C in SF-900 II media supplemented with 1.5 % FCS and 2 mM L-Glu were infected at $10^6$ cells/mL and an MOI of 0.1 in a 0.5 L CELLSPIN Spinner. After 72 h, cells were collected by centrifugation at 300 ×g for 15 mn, washed in PBS and stored at -20°C.

***Homology modelling***

The crystal structure of Osh1 ORD (PDB: 5H2D) (5) was used as template to model human OSBP ORD. The sequence alignment was improved using psi-blast web tools (6). One hundred models (with for each of these models ten loops) of OSBP ORD were constructed separately by comparative modelling using the MODELLER tool (version 9.12) (7). The five best models with the lowest value of the MODELLER objective function were selected. The stereo-chemical quality of each model was subsequently evaluated using the SAVES web tool (http://servicesn.mbi.ucla.edu/SAVES/). The best model was finally subjected to a steepest descent energy minimization with GROMACS tool 2018 version (8) using CHARMM36 force field (9). For this, a cubic box solvated with Tip3p water molecules and 120 mM NaCl was built to neutralize the system.

***LC-MS***
RPE-1 cells were treated with drugs during 2 h in complete growth medium at 37°C, washed once with ice-cold PBS and then scratched in 1 mL PBS (4°C). Cells were then centrifuged, resuspended in $H_2O$ at 4°C and centrifuged. Pellets were dried under argon and frozen. Lipids from cell pellets were extracted according to a modified Bligh and Dyer protocol (10). Cell pellets were resuspended in 200 µL $H_2O$ and transferred in a glass tube containing 500 µL of methanol and 250 µL of chloroform. The mixture was vortexed for 30 s and centrifuged (2500 rpm, 4°C, 10 min). 300 µL of the organic phase was collected in a new glass tube and dried under a stream of nitrogen. The dried extract was resuspended in 60 µL of methanol/chloroform 1:1 (v/v) and transferred in an injection vial before liquid chromatography and mass spectrometry analysis. Reverse phase liquid chromatography was selected for separation with an UPLC system (Ultimate 3000, ThermoFisher). Lipid extracts were separated on an Accucore C18 150x2.1, 2.5µm column (ThermoFisher) operated at 400 µl/min flow rate. The injection volume was 3 µL of diluted lipid extract. Eluent solutions were ACN/$H_2O$ 50/50 (V/V) containing 10 mM ammonium formate and 0.1% formic acid (solvent A) and IPA/ACN/$H_2O$ 88/10/2 (V/V) containing 2 mM ammonium formate and 0.02% formic acid (solvent B). The used step gradient of elution was: 0 min 35% B, 0.0-4.0 min 35 to 60% B, 4.0-8.0 min 60 to 70% B, 8.0-16.0 min 70 to 85% B, 16.0-25 min 85 to 97% B, 25-25.1 min 97 to 100% B, 25.1-31 min 100% B and finally the column was reconditioned at 35% B for 4 min.
The UPLC system was coupled with a Q-exactive orbitrap Mass Spectrometer (Thermofisher, CA); equipped with a heated electrospray ionization (HESI) probe. This spectrometer was controlled by the Xcalibur software and was operated in electrospray positive mode. MS spectra were acquired at a resolution of 70 000 (200 m/z) in a mass range of 250−1200 m/z. 15 most intense precursor ions were selected and isolated with a window of 1 m/z and fragmented by HCD (Higher energy C-Trap Dissociation) with normalized collision energy (NCE) of 25 and 30 eV. MS/MS spectra were acquired in the ion trap in a mass range of 200-2000 m/z, the resolution was set at 35 000 at 200 m/z. Data were reprocessed using Skyline 4.1.0.18169 with transition list settings including precursor m/z and precursor charge.
For lipid mass analysis from OSBP-ORD, the purified protein was first dialyzed in Tris 2 mM, NaCl 300 mM, and lipids were extracted according to a modified Bligh and Dyer protocol (10), and separated by chromatography on a C18 column as described above. For quantification purpose, lipids were extracted from a known amount of ORD (2.8 nmoles) together with ergosterol (2.5 nmoles), which was used as an internal standard. The extraction yields of cholesterol and ergosterol were assumed to be similar. In parallel, mass analysis of 0, 1, 2.5, 5 and 10 nmol cholesterol and ergosterol standards was performed to establish the differential ionization yield, so as to calculate the cholesterol/ORD ratio.

**Figure S1. Effect of OSBP silencing on TGN-to-plasma membrane trafficking.**
**A.** Time-lapse imaging of RPE-1 cells stably expressing SBP-EGFP-CD59, treated either with control siRNA (siNT) or siRNA against OSBP for 48 h, and incubated with biotin as indicated. Real-time images were acquired using a wide-field microscope at 2 frames/min. Scale bars: 20 µm. **B.** Normalized intensity of SBP-EGFP-CD59 at the Golgi. Data are means ±SEM (n=6).

**Figure S2. SWG stabilizes ER-TGN contacts mediated by the OSBP/VAP-A complex.**
**A.** RPE-1 coexpressing TagBFP-βGalT1, GFP-VAP-A, and mCherry-OSBP. βGalT1 labels the TGN and VAP-A the ER network, whereas OSBP is mostly cytosolic. Upon SWG treatment (10 nM, at 37°C), OSBP and VAP-A concentrate to the perinuclear region. Scale bar: 20 µm. **B.** Normalized fluorescence intensities at the Golgi area over time. Data are means ±SEM (n=3).

**Figure S3. SWG rapidly crosses the plasma membrane and accumulates in OSBP-enriched regions.**
**A.** Epifluorescence images of RPE-1 cells labelled with SWG (1 µM) for the indicated time in growth medium at 37°C, fixed, permeabilized and processed for immunofluorescence to assess the localization of endogenous OSBP and TGN-46. Scale bar: 20 µm. **B.** Fluorescence intensity at the TGN of the Alexa Fluor-conjugated antibody directed against primary anti-OSBP over time. Data are means ± SEM (n=10). **C.** Time-lapse epifluorescence images of RPE1-derived GPMVs labelled with WGA-Alexa 488 upon addition of SWG (5 µM). The bar corresponding to the line scan: 16 µm. **D.** SWG fluorescence intensity measured over time at plasma membrane and in cytosol regions (the line scan intensity was used to determine “membrane” intensity, which was the mean area under the two peaks, and the “cytosol” intensity, which was the average lumenal intensity). Data are means ±SEM (n=3).

**Figure S4. SWG loading and lipid binding status of OSBP-ORD.**
**A.** Same experiment as presented in Figure 4F and 4G (thesit 0.1%) (Zoom on the first time points). **B.** Lipid mass analysis from purified OSBP-ORD. Cholesterol was evidenced in the preparation by LC-MS. Quantitative analysis indicated that the cholesterol/ORD ratio is of about 0.65, suggesting that a majority of the ORD is purified with a cholesterol molecule in the lipid-binding cavity.

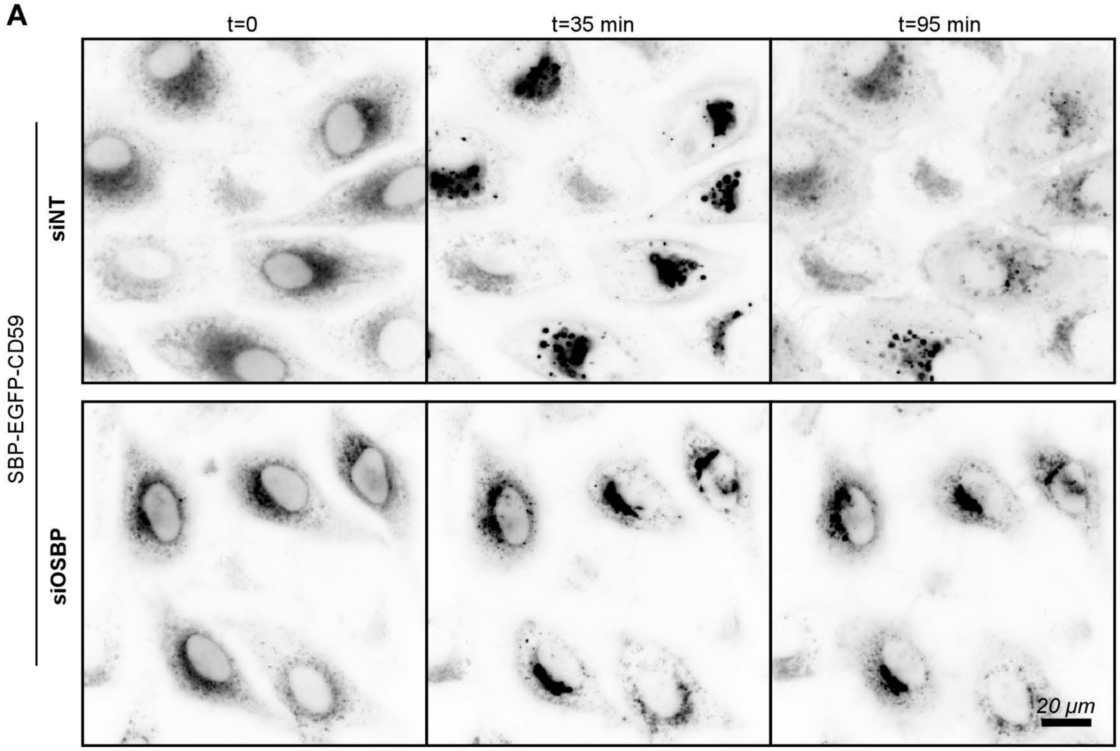
A
t=0
t=35 min
t=95 min
SBP-EGFP-CD59
siNT
siOSBP
20 µm

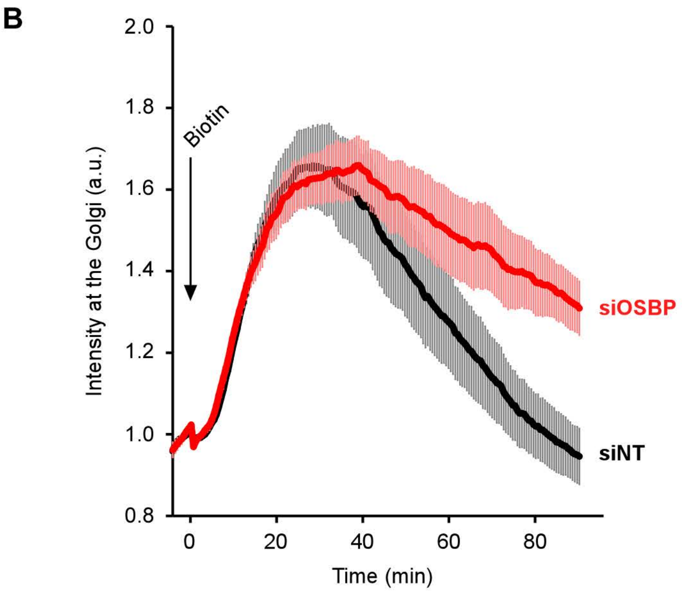
B
Biotin
Intensity at the Golgi (a.u.)
2.0
1.8
1.6
1.4
1.2
1.0
0.8
0
20
40
60
80
Time (min)
siOSBP
siNT

Figure S1

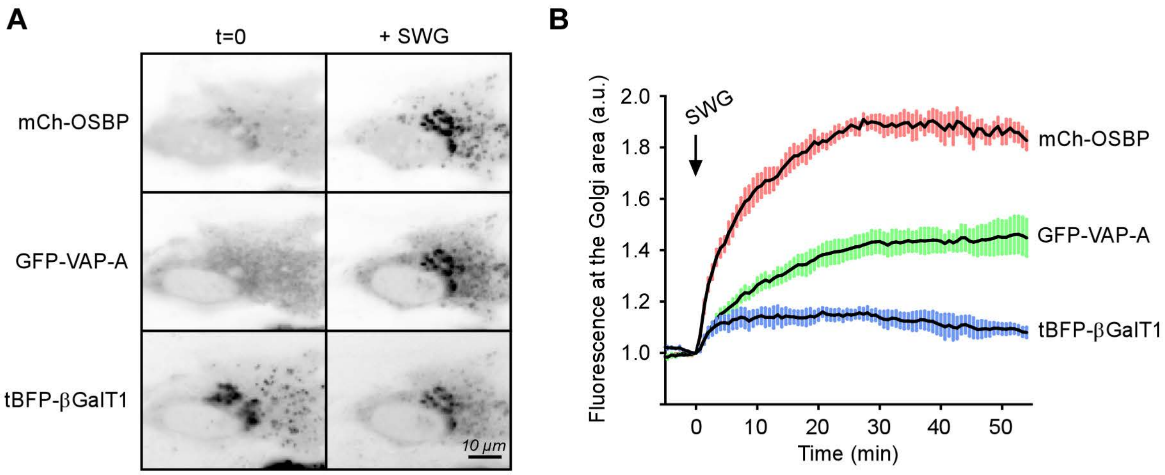
A
t=0
+ SWG
mCh-OSBP
GFP-VAP-A
tBFP-βGalT1
10 µm
B
SWG
2.0
1.8
1.6
1.4
1.2
1.0
0
10
20
30
40
50
Fluorescence at the Golgi area (a.u.)
Time (min)
mCh-OSBP
GFP-VAP-A
tBFP-βGalT1

Figure S2

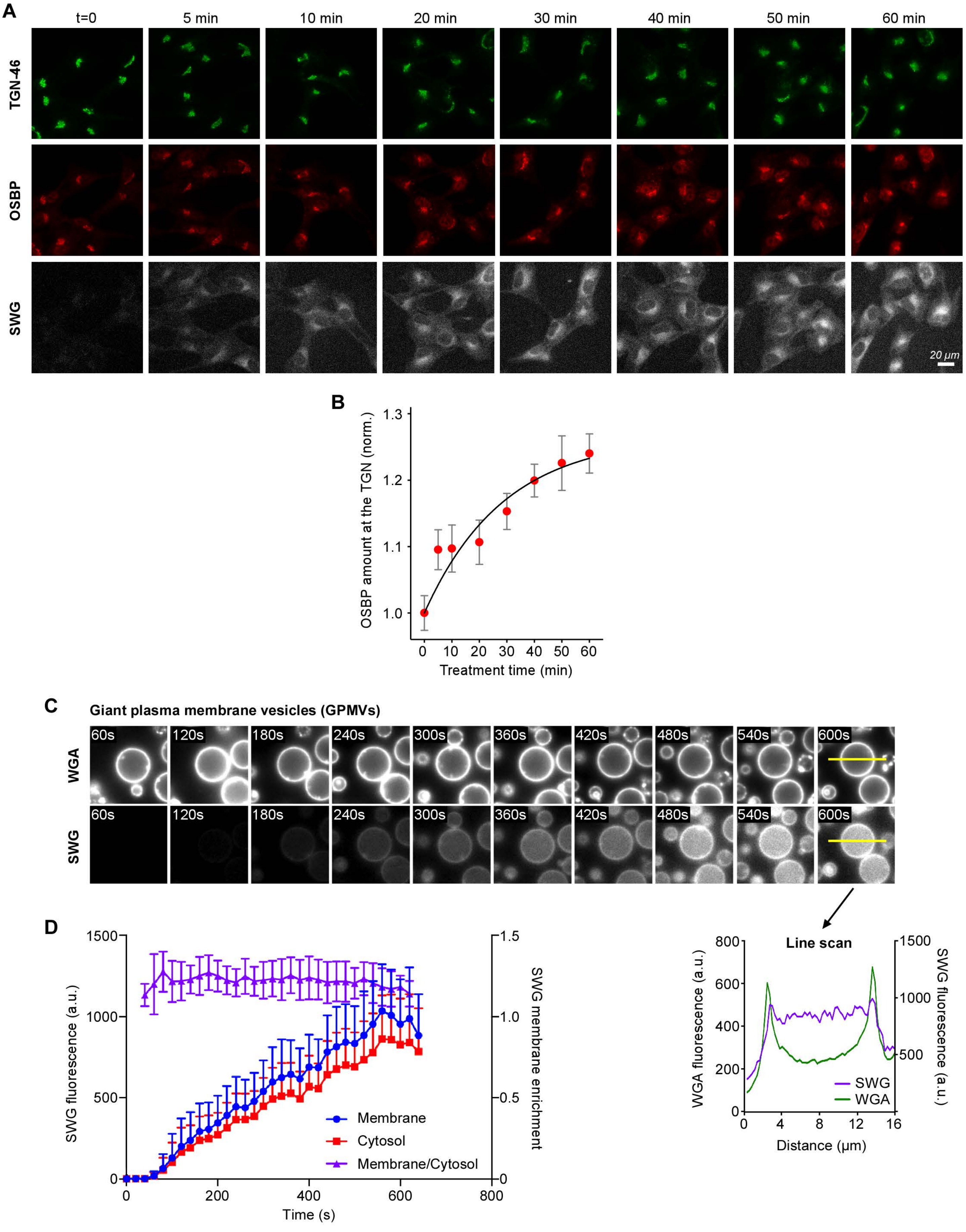
A
t=0
5 min
10 min
20 min
30 min
40 min
50 min
60 min
TGN-46
OSBP
SWG
20 µm
B
OSBP amount at the TGN (norm.)
1.0
1.1
1.2
1.3
0
10
20
30
40
50
60
Treatment time (min)
C
Giant plasma membrane vesicles (GPMVs)
WGA
SWG
60s
120s
180s
240s
300s
360s
420s
480s
540s
600s
D
SWG fluorescence (a.u.)
0
500
1000
1500
SWG membrane enrichment
0
0.5
1.0
1.5
Membrane
Cytosol
Membrane/Cytosol
0
200
400
600
800
Time (s)
Line scan
WGA fluorescence (a.u.)
0
200
400
600
800
SWG fluorescence (a.u.)
500
1000
1500
SWG
WGA
0
4
8
12
16
Distance (µm)

Figure S3

**A**

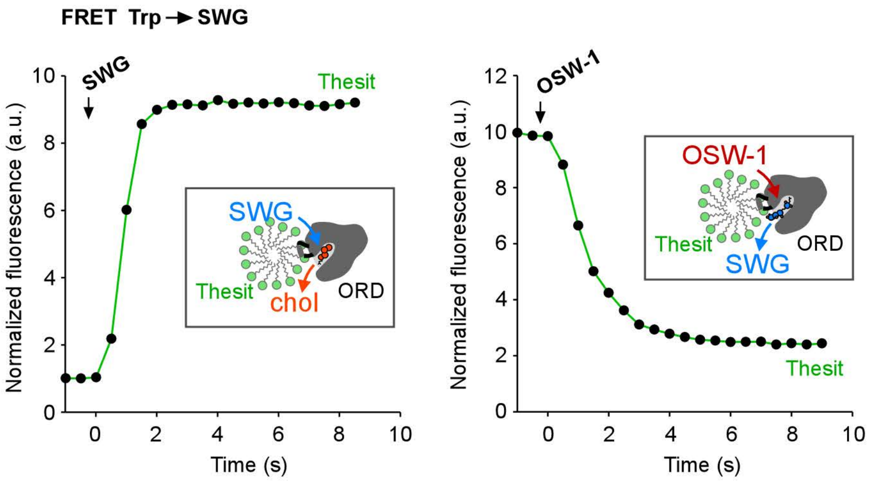
FRET Trp → SWG
SWG
Thesit
SWG
Thesit
chol
ORD
Normalized fluorescence (a.u.)
Time (s)
OSW-1
OSW-1
Thesit
ORD
SWG
Thesit
Normalized fluorescence (a.u.)
Time (s)

**B**

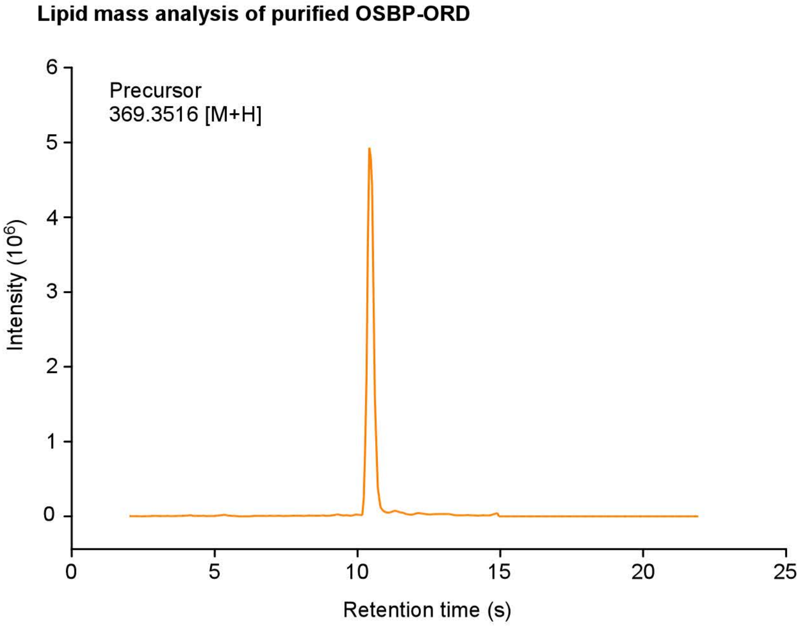
Lipid mass analysis of purified OSBP-ORD
Precursor
369.3516 [M+H]
Intensity (10^6)
Retention time (s)

Figure S4
